# The genetic basis of variation immune defense against *Lysinibacillus fusiformis* infection in *Drosophila melanogaster*

**DOI:** 10.1101/2022.10.19.512815

**Authors:** Brittny R. Smith, Kistie B. Patch, Robert L. Unckless

## Abstract

The genetic causes of phenotypic variation often differ depending on the population examined, particularly if the populations were founded by relatively small numbers of genotypes. Similarly, the genetic causes of phenotypic variation among similar traits (resistance to different xenobiotic compounds or pathogens) may also be completely different or only partially overlapping. Differences in genetic causes for variation in the same trait among populations suggests considerable context dependence for how selection might act on those traits. Similarities in the genetic causes of variation for different traits, on the other hand, suggests pleiotropy which also would influence how natural selection would shape variation in a trait. We characterized immune defense against a natural *Drosophila* pathogen, the Gram-positive *Lysinibacillus fusiformis*, in three different populations and found almost no overlap in the genetic architecture of variation in survival post infection. However, when comparing our results to a similar experiment with the fungal pathogen, *B. bassiana*, we found a convincing shared QTL peak for both pathogens. This peak contains the *Bomanin* cluster of *Drosophila* immune effectors. RNAi knockdown experiments confirms a role of some of these genes in immune defense against both pathogens. This suggests that natural selection may act on the entire cluster of *Bomanin* genes (and the linked region under the QTL) or specific peptides for specific pathogens.

**Author Summary:** Like most traits, the way individuals respond to infection vary among individuals within a population. Some of this variation is caused by genetic differences in the host organism. Over the past decade, two prominent resources were developed to assess genetic variation for complex traits of the fruit fly, *Drosophila melanogaster* and map the genetic variants responsible. We recently described a strain of *Lysinibacillus fusiformis* bacteria, which was isolated from fruit flies and is moderately virulent when flies are infected. We mapped genetic variation in resistance *L. fusiformis* using these mapping resources. We find that among the resources, different changes were associated with immune defense. However, we also found that within a resource, the same region of the genome was associated with resistance to both *L. fusiformis* and a fungal pathogen. These results suggest that different populations adapt differently to the same pathogens, but two distinct pathogens share similar causes of genetic variation within a single population.

## Introduction

Infection is ubiquitous, but the ways in which different organisms fight infection with different pathogens is likely as varied as the organisms themselves. Within species, there is often enormous genetic variation in the ability to survive infection and it is not clear whether the same host variants will provide wide ranging protection against several pathogens or whether specific host alleles provide narrow protection against a few pathogens [1, 4-10]. If certain host alleles are indeed generally better at fighting infection, it follows that those alleles must either be in the midst of sweeping through the population *or* that their fitness benefits are context dependent. Perhaps alleles conferring resistance are energetically more costly or cause dysbiosis of the native microbiota [11-13].

*Drosophila* and other invertebrates lack an adaptive immune system, which allows us to focus on how the innate immune system functions in infection and disease [14]. As a result, some of the initial insight about immune defense pathways were gained by studying *Drosophila* and such advances continue today [14-17]. Pioneering work in the 1980s provided insight into the Toll and Imd pathways which are largely conserved in vertebrates [18]. Recently, we’ve worked out more specific associations between particular immune effectors and infection outcomes [5, 19, 20]. In addition to the functional genetic work on the immune system, others have focused on the role of natural genetic variation in resistance to infection [1, 21, 22]. In some cases, those studies have inspired a deeper functional genetic dissection of the roles of specific immune genes in infection [23].

Insect innate immunity can be broadly grouped into cellular and humoral immunity. Cellular immunity involves the differentiation of blood cells that have several roles in directly attacking and/or immobilizing pathogens [24]. In contrast, the humoral immune response involves signaling cascades that generally lead to enormous induction of immune effectors [25]. In *Drosophila*, the canonical view is that the Toll pathway is activated in response to Gram-positive and fungal infection and stimulates expression of immune and antimicrobial peptide genes (*Bomanins, Daishos, Drosomycins, etc*.)[14, 26, 27]. This canonical view, suggests that the Imd pathway is activated in response to Gram-negative infection and stimulates the expression of antimicrobial peptides (*Diptericins, Cecropins, Defensins, Attacins, etc*.). Other pathways including JNK and JAK/STAT also play important roles and likely interact with Toll and Imd [28, 29].

Dissecting the genetic bases of phenotypic variation in a phenotype such as immune defense yields two important results. First, because the associated approaches are unbiased, they often reveal genes previously unknown to be involved in immune pathways. Second, an understanding of the genetic causes of phenotypic variation informs both genetics and evolution. There are two commonly used *D. melanogaster* panels used for mapping genetic variation in phenotypes. The first is the Drosophila Genetic Reference Panel (DGRP) which was derived from a set of gravid females collected at a farmer’s market in North Carolina, USA [30]. The offspring of those gravid females were inbred for several generations, then 205 lines were sequenced and made available to the public. This allows for a standard genome-wide association study (GWAS) approach to determining the genetic basis of phenotypic variation and has been used widely for several different phenotypes [31-36], including response to infection [21, 37, 38]. The second panel is the Drosophila Synthetic Population Resource (DSPR) [39, 40]. This set of lines (actually two mostly independent sets) were derived from eight inbred founder lines that were allowed to interbreed for several generations, then were inbred. Founders were sequenced to high depth and the recombinant inbred lines (RILs) were genotyped at low depth to assign founder genotypes across the genome. This approach allows a quantitative trait locus (QTL) approach [41, 42]. The DSPR lines have not been used for studies of genetic variation in infection, except for the fungal insect pathogen *Beauvaria bassiana* [43] and the *Drosophila* C Virus [44].

Our study investigated natural variation in defense against the Gram-positive *Lysinibacillus fusiformis*, which was isolated from *Drosophila* in the lab [45]. We measured survival 2- and 5-days after septic injury with *L. fusiformis* in the DGRP population and both DSPR populations (A and B). We also performed RNA-sequencing of flies 12-hours post infection and found that induction of gene expression after infection with the Gram-positive *L. fusiformis* is better correlated with induction of gene expression after Gram-negative microbes than with other Gram-positive microbes. This may be because the peptidoglycan, which is responsible in part for immune induction, of *L. fusiformis* is *not* similar to that of most Gram-positive microbes. Though the DGRP analysis revealed an excellent candidate gene, we found very little overlap in significantly associated SNPs with other immune-related studies. In contrast, the DSPR A population has a QTL peak that overlaps with a single peak in the *B. bassiana* study. While this peak contains several genes, we explored the possibility that variation in the same genes is responsible for infection differences for both *L. fusiformis* and *B. bassiana*.

## Results

### L. fusiformis is moderately virulent and leads to induction of genes in volved in immune defense

We infected males from the standard *D melanogaster* lab line, Canton S, with *L. fusiformis* at an optical density (OD_600_) of 3.977 and measured survival for seven days post infection (DPI). At this dose, 76% of flies (51 of 67) survived to 7 days, while all flies pricked with a sterile needle survived (Figure 1A). This represents a significant, but moderate reduction in survival (Cox Proportional Hazard test, hazard ratio = 16.1, P=0.007). Note that to achieve sensible hazard ratio, we called one sterile pricked fly dead at day 7. At this point, we could have elected to use a higher dose for subsequent infection experiments, but the OD_600_ around 4.0 led to considerable variation in survival among different lines (see below), so we continued using this dosage throughout the remainder of the experiment.

**Figure 1.**
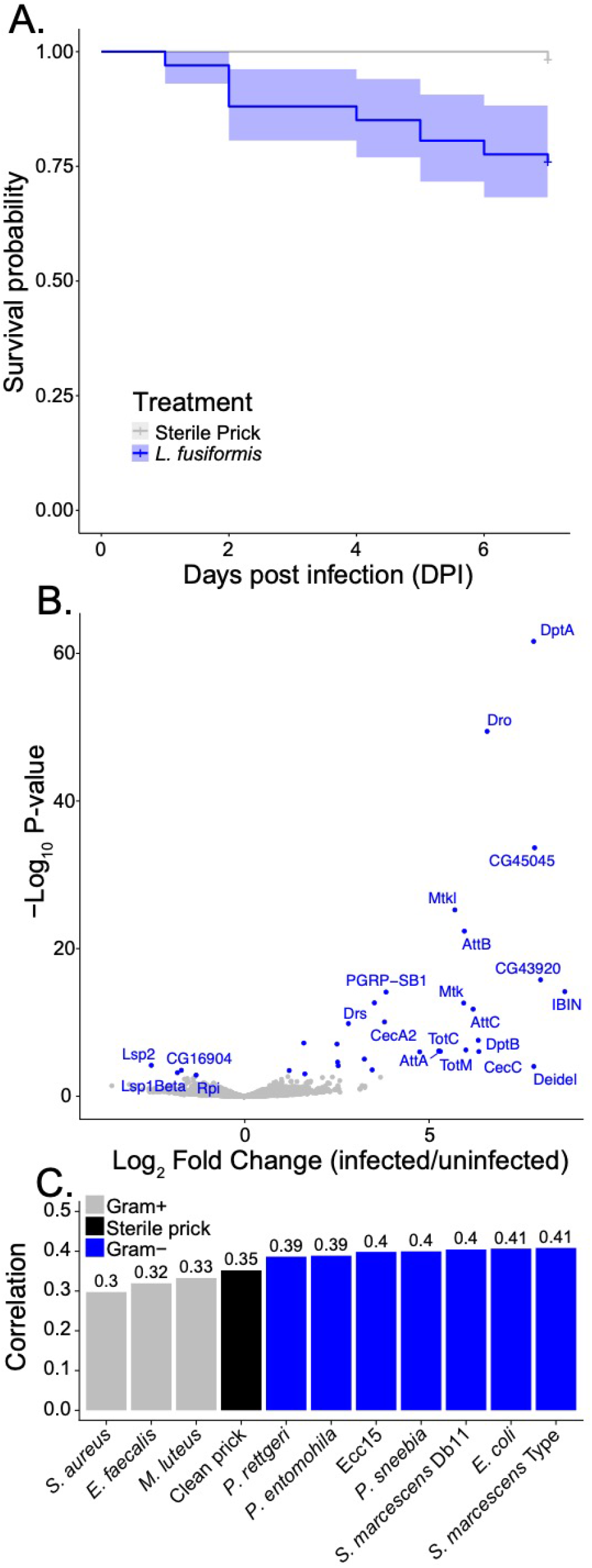
*Lysinibacillus fusiformis* infection is moderately virulent. A) Canton S male survival after infect with *L. fusiformis* (OD_600_=3.977, blue, n=67) or sterile prick (gray, n=59) with 95% confidence intervals. B) Volcano plot of gene expression in Canton S males 12 hours post infection with OD_600_=3.977 *L. fusiformis* vs. uninfected (no sterile prick). Blue points are significantly differentially expressed (FDR-adjusted P-value<0.05) and genes of interest are labeled. C) Pearson correlation coefficients for Log_2_ fold change between infections with *L. fusiformis* and microbes from Troha *et al*. 2018.

We performed RNA-sequencing of pools of males 12 hours post-infection with *L. fusiformis* to examine changes in gene expression upon infection (Figure 1B). The most induced genes are expected based on other studies [46-48] and are mostly antimicrobial peptides in the Toll and Imd pathways. We used an adjusted P-value threshold of 0.1 for Gene Ontology enrichment analysis, which yielded 36 genes differentially expressed. Gene Ontology categories significantly enriched included several related to immune defense: response to bacteria, response to external biotic stress, defense response, immune response, defense response to Gram-positive bacterium, defense response to Gram-negative bacterium (see the entire list in Supplemental Table 1).

**Table 1.**
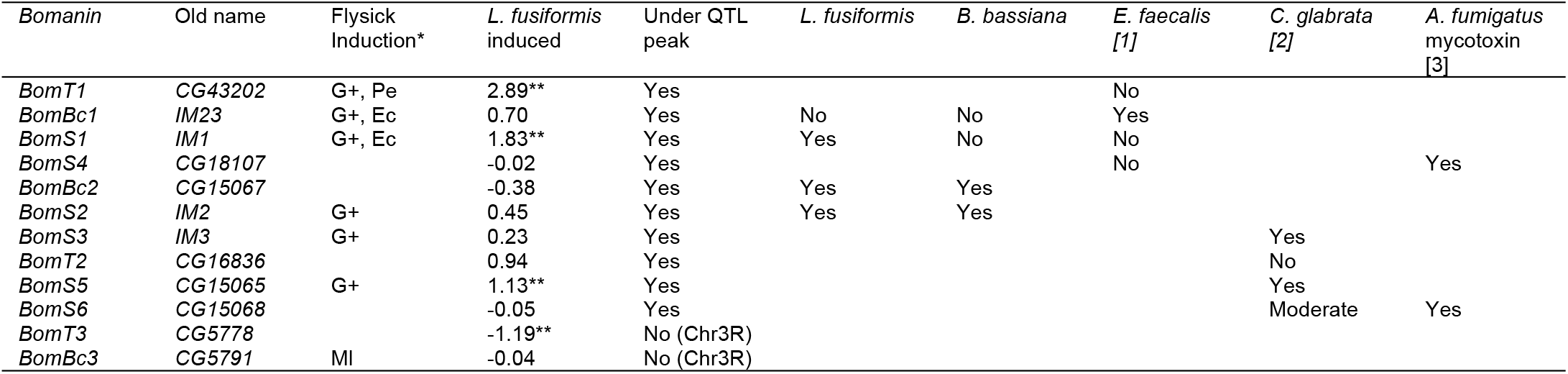
Summary of *Bomanin* gene immune phenotypes. *Flysick induction indicates which of the ten microbes resulted in at least 10-fold higher expression compared to uninfected individuals. “G+” means all Gram-positive bacteria (*M. luteus - Ml, E. faecalis, S. aureus*) induced at least 10-fold. Two Gram-negative bacteria out of 7 also led to a 10-fold increase in expression: *Pseudomonas entomophila* (Pe) and *Escherichia coli* (Ec).

It appears that the induced antimicrobial peptides are more squarely induced by the Imd pathway instead of the Toll pathway, suggesting induction of gene expression is more similar to Gram-negative bacteria than Gram-positive bacteria. The *Lysinibacillus* genus are Gram-positive bacteria, but like others in the Bacillales order, the peptidoglycan is not l-lysine-(lys) type. Instead, most *Lysinibacillus* examined have the A4alpha (Lys-Asp) peptidoglycan [49, 50]. Since the humoral response in *D. melanogaster* is driven in part by recognition of peptidoglycan type, the expected relative responses of the Toll and Imd responses to Bacillales is less clear cut than for most bacteria that fit more neatly into Gram+ and Gram-categories based on peptidoglycan [14]. We calculated Pearson correlation coefficients for Log_2_ Fold Change (infected vs. uninfected) between our *L. fusiformis* data and 12-hour post infection gene expression for 11 bacterial infection treatments from the Troha *et al. [46]* data. Although all correlations were significant (P<0.0001), there was stronger correlation between *L. fusiformis* and Gram-negative bacteria (median = 0.4) than any of the Gram-positive bacteria (median = 0.32, Figure 1C). The same pattern is true when we only considered significantly differentially expressed genes or only significantly induced genes (data not shown). Thus, *L. fusiformis* is a moderately virulent bacterial pathogen that induces a strong innate immune response that is more similar to the transcriptional response to Gram-negative than Gram-positive bacteria.

### Genetic variation for immune defense against L. fusiformis in the DGRP

All three populations (DGRP and two DSPR populations) showed considerable genetic variation for survival after infection with *L. fusiformis* (Figures 2A, 3A and 4A). Raw survival in the DGRP population ranged from 10% to 100% surviving 5 days post infection (DPI). We also examined survival 2 days post infection, which was much higher overall and showed less variation (Supplemental Figure 1). Like several other studies of immune defense in the DGRP, we found that line 321 was particularly susceptible to infection, so we calculated line effects with and without line 321 (Supplemental Table 2, Supplemental Figure 2). The heritability in the DGRP was 0.802 for survival 5 DPI (0.759 if line 321 is removed) and 0.924 for survival 2 DPI (0.890 if line 321 is removed).

**Figure 2:**
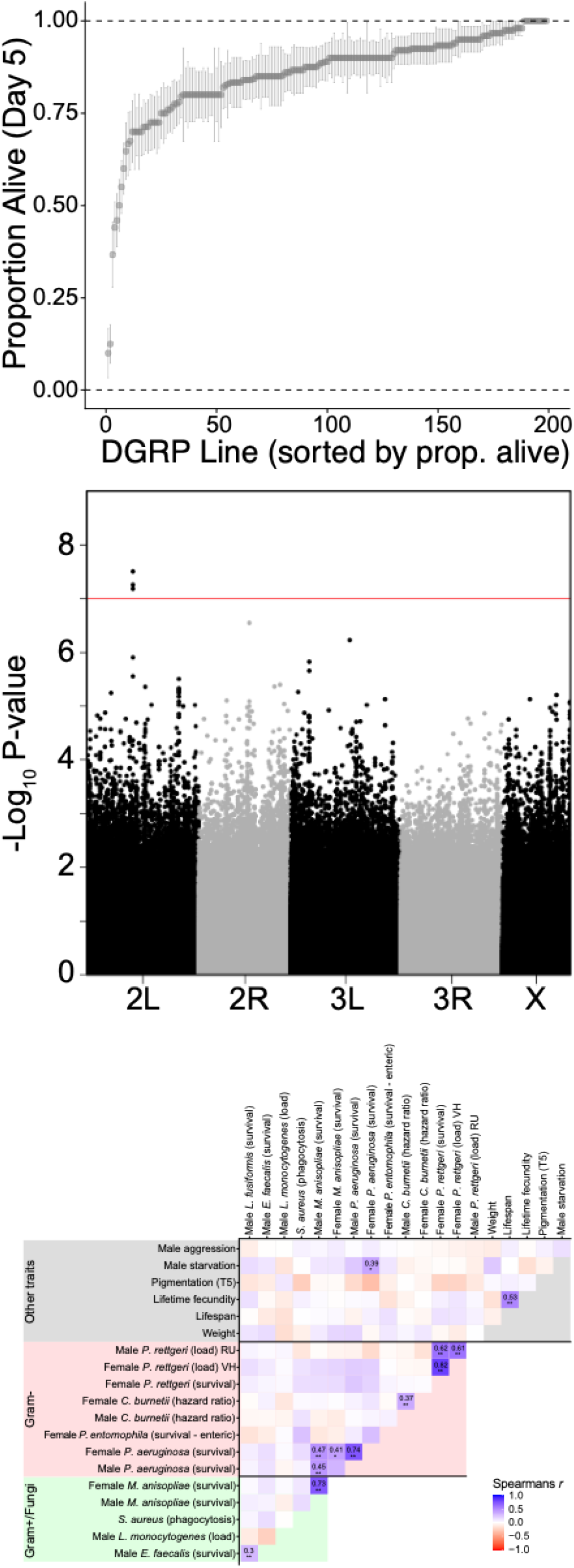
Genetic variation in immune defense against *L. fusiformis* in the DGRP. A) Sorted raw line mean proportion alive 5 days post infection with standard error of the proportion. B) Manhattan plot of genome wide associations with 5-day survival random effects. C) Correlation of line effects from several DGRP immune studies.

**Figure 3.**
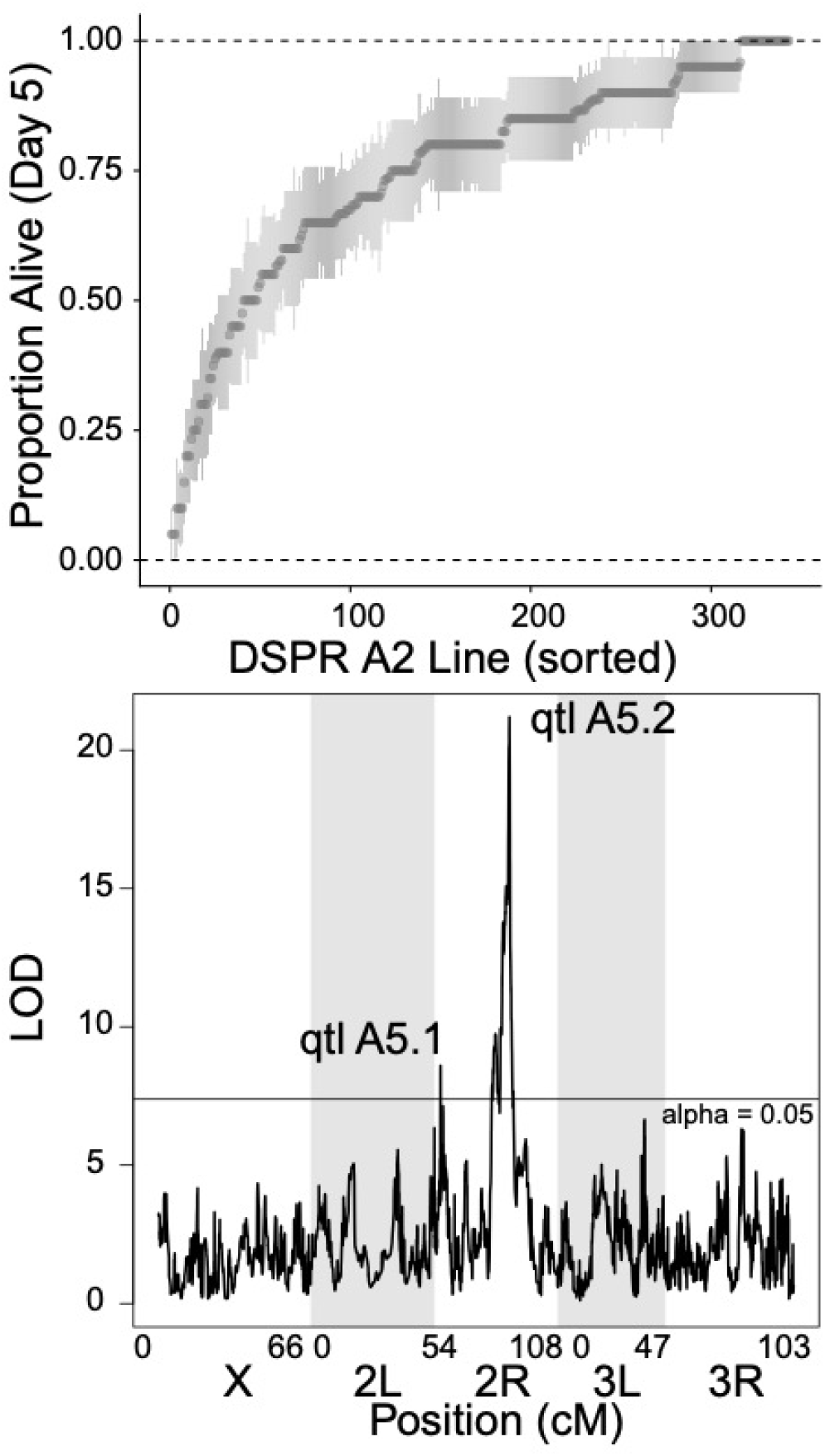
Genetic architecture of immune defense against *L. fusiformis* in DSPR population A2. A) Raw line means for proportion alive at day 5 post infection with standard error of the proportion sorted be mean proportion. B) QTL plot for line effects of survival post infection with *L. fusiformis*. Two QTLs are labeled, horizontal line shows a significance threshold with alpha = 0.05.

**Figure 4.**
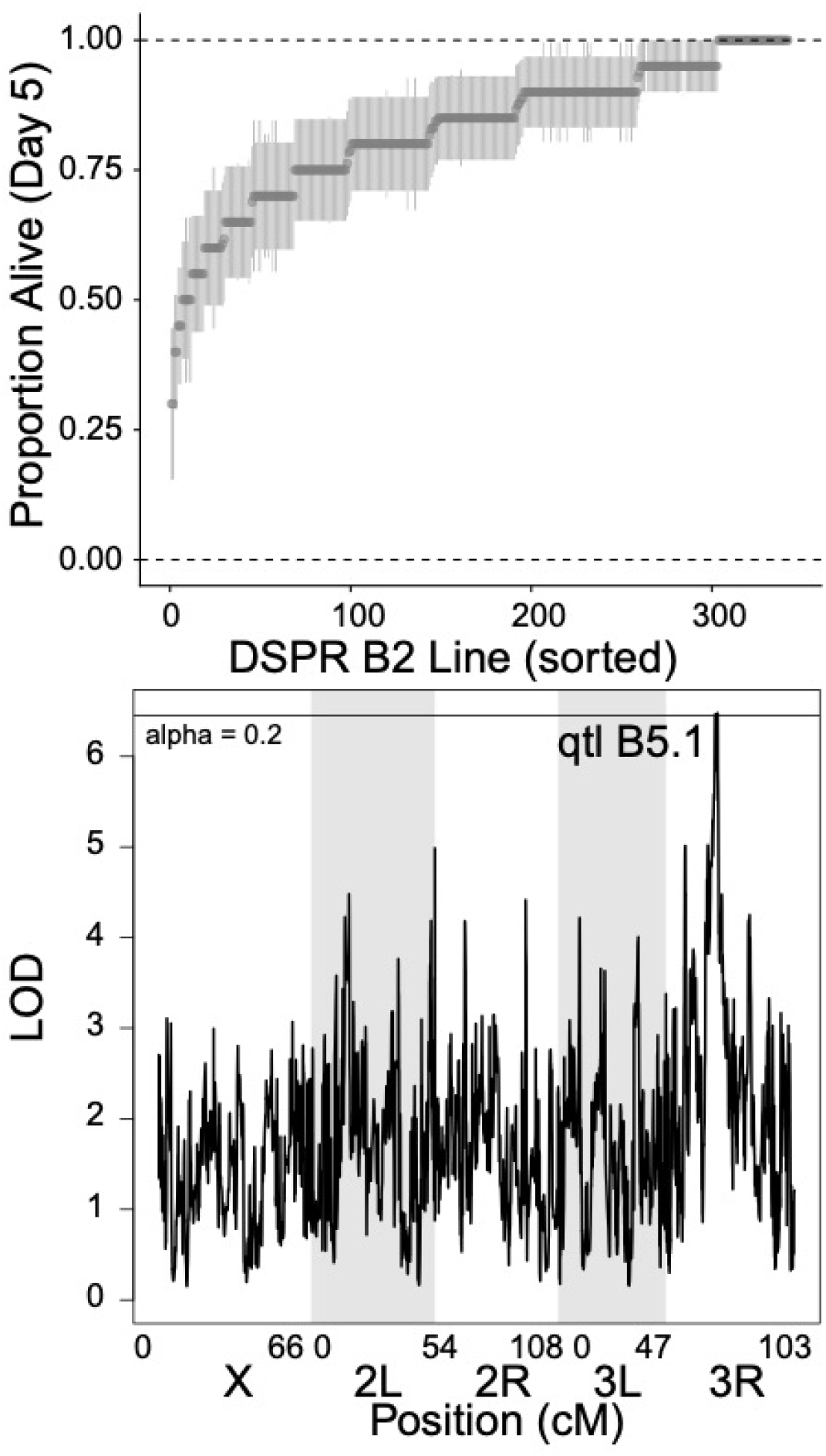
Genetic architecture of immune defense against *L. fusiformis* in DSPR population B2. A) Raw line means for proportion alive at day 5 post infection with standard error of the proportion sorted be mean proportion. B) QTL plot for line effects of survival post infection with *L. fusiformis*. Two QTLs are labeled, horizontal line shows a significance threshold with alpha = 0.2.

We used the online DGRP mapping tool because it accounts for inversions, *Wolbachia* infection status and the relatedness of lines. We performed the mapping for 5-day survival and 2-day survival both with and without line 321). We chose an FDR cutoff of 10% as our significance threshold, and few SNPs were significant at that cutoff (Figure 2B, Supplemental Figure 3, Supplemental Tables 3-6). Below we discuss the different mapping approaches, but put the most faith in mapping results of 5-day survival when we discarded line 321 because those results provided the most reasonable QQ plot, an indication that the mapping model was reasonable (Supplemental Figure 4).

The FDR-adjusted significance threshold of 0.1 yielded only a handful of significant SNPs (Supplemental Tables 3-6). Sixteen SNPs were significantly associated with survival 5 DPI when all lines were considered. Eleven intronic SNPs were in *eys*/*CG9967, Pvr, CG44153, Trim9, CG15611* (4 SNPs), *trio, CG32698* (actually a 2bp deletion), and *Frq1*. One SNP was in the exon of the noncoding RNA *CR44310*, one SNP caused a silent change in *CG14971*, and 3 SNPs were intergenic. When we omitted line 321, only 3 SNPs were significantly associated with 5-day survival – all three were intronic SNPs in *Pvr*. Note that the SNPs significantly associated when all lines were included but not significantly associated when 321 was omitted were mostly (13 out of 14) still tested, but had low enough minor allele frequencies that power was lost and P-values were above our significance threshold. Only one fell below our minor allele frequency threshold and so was not tested. Eight SNPs were significantly associated with 2-day survival when all lines were considered. This included 4 intronic SNPs in *Zasp52, trio, CG32364*, and *Plod* as well as 4 SNPs in the 3’ UTR of *CG15564*. There were no significantly associated SNPs 2-days post infection when line 321 was omitted. The SNPs in *Pvr* significantly associated at 5-days post infection were nominally significant at 2-days post infection (∼3.0 × 10^−4^ when all lines were considered and ∼4.0 × 10^−5^ when line 321 was omitted).

We investigated genes connected to significantly associated SNPs in a few ways. First, we examined whether any of these genes were differentially expressed 12 hours after infection based on our RNA-seq data. None were significantly differentially expressed even if we consider the nominal P-values, so genes connected to significantly associated SNPs are not significantly induced or repressed upon infection (at least at the 12-hour point).

Next, we examined annotated functions of these genes. Two stand out. First, *Pvr* (*PDGF- and VEGF-receptor related*) is involved in the regulation of several innate immune functions including apoptosis, humoral immune response, JNK signaling and wound healing [51-55]. *Pvr* is a negative regulator of humoral immune response, so if these SNPs are associated with alternative splicing or otherwise influence *Pvr* expression, that would be a plausible connection. The other gene of interest is *CG44153*, which is predicted to enable cell-cell adhesion mediator activity and is active in the plasma membrane [56]. It stands out because it was also identified in two other DGRP immunity association studies: survival time after infection with *Pseudomonas aeruginosa [21]*, and microbiota-related diet shift [57]. However, *CG44153* also was reported to harbor genetic variation with several other non-immune phenotypes, so it is possible that there is something about genetic variation at the locus that makes it prone to false positive associations.

### Little evidence for correlations among immune phenotypes in the DGRP

The DGRP lines have been available to the public for more than a decade, and in that time, there have been several studies attempting to map variation in immune defense against a range of pathogens. This allows us to begin to determine whether immune defense against different pathogens is correlated. Are lines that are better at fighting infection against one pathogen, generally better at fighting infection? If so, is this true universally or within categories such as Gram-positive bacteria? A general correlation in immune defense would suggest that at least some of the genetic basis of that variation is also shared for these different pathogens. There is some evidence for this in the overlap in associations in this study and those of Chapman *et al*. and Wang *et al*. [1, 21]. However, it is also possible that resistance is much more specific for each pathogen. This is likely given some of the specificity of associations – particularly for *Providencia rettgeri* and *Diptericin* [5, 20, 37].

We calculated Spearman’s rank correlations for 14 DGRP immune-related data sets plus an additional 6 unrelated phenotypes (Figure 2C). Though there are several positive correlations, few are significantly correlated after Benjamini & Hochberg correction. Correlations tend to be between the same microbe using different experimental design (different sex or survival vs. bacterial load) or between different microbes measured in the same laboratory (*E. faecalis* and *L. fusiformis* were both measured in our laboratory at the same basic time; *M. anisopliae* and *P. aeruginosa* were reported as part of the same experiment in Wang et al. [21]). However, there is an almost complete lack of significant correlation between immune phenotypes and any of the other phenotypes, the exception being a positive correlation between female survival after *P. aeruginosa* and male starvation survival. Given the general lack of correlation between phenotypes, we decided not to look at the overlap in significantly associated SNPs between immune phenotypes. Our correlation results suggest that the DGRP immune phenotypes that have been reported do show correlation, but it is difficult with the available data to determine if these correlations are due to phylogenetic relatedness of the microbes used or are an artifact of something about the experimental design (how phenotypes were defined, dose, etc.) or labs where the experiments were performed.

### Genetic variation for immune defense against L. fusiformis in the DSPR

The DSPR consists of two populations (A and B) that were derived from eight different founder genotypes. Each of those populations was further split into two parallel sets (*e*.*g*. A1 and A2) which were derived from the same founders, but different interbreeding populations. We infected 343 lines from population A2 with *L. fusiformis* (OD_600_ ∼ 4.0, mean = 22.3 male flies per line, median = 20 male flies per line, standard deviation = 6.47 male flies per line). The overall survival rate 5 days post infection was 74.76% and the line mean raw survival was 75.16% (median = 80%, standard deviation = 21.11%, Figure 3A, Figure S5A for Day 2). Heritability was 0.828 for survival 5 DPI and 0.928 2 DPI.

We used the same logistic regression model as described above to calculate line effects for each DSPR population A line (Supplemental Figure 6A, Supplemental Table S7), and used the DSPRqtl package [58] to determine regions of the genome associated with variation in immune defense against *L. fusiformis*. We determined a significance threshold (alpha = 0.05) by permuting the data for 1000 iterations. This yielded several QTL peaks with LOD scores above the threshold, but they concentrate in two regions (Figure 3B). The first peak (qtl A5.1) was centered at Chr2R:3,420,000 (LOD drop 2 range 3,360,000 to 3,470,000, note this is release 5 of the *D. melanogaster* genome, LOD = 8.59) and was associated with 10.96% of the variation in the phenotype. We extracted the effects of individual founder genotypes (Supplemental Figure 7A) and found that founder A1 genotypes at this region had significantly higher survival than the other four genotypes represented. This includes all or part of 21 genes, none of which have obvious immune function, nor were any differentially expressed in our RNA-seq data after p-value correction. One gene, *Cyt-b5* showed an almost 50% reduction in expression upon infection, which was associated with a nominal p-value of 0.02. Importantly it is annotated to be involved in metabolism of xenobiotic compound, but the naïve expectation if it is breaking down microbial toxins would be induction, not suppression upon infection.

The second peak (qtl A5.2) had a higher LOD score and was also much broader. We focused on the main peak (highest LOD score) which was centered at Chr2R:14,720,000 (LOD drop range from 14,680,000 to 14,770,000, LOD = 21.19) and was associated with 24.89% of the variation. This peak contains all or part of 19 genes including two genes involved in immune defense: *DptA* and *DptB*. The founder genotype from A5 is associated with significantly lower survival than the other represented genotypes (Supplemental Figure 7B). If we expand to look at genes under the three adjacent (actually overlapping) peaks with LOD scores above 20, we also include the gene, *Jabba*, which also functions in immune defense. One more QTL peak still overlapping these but with a LOD score of 14.7 (peak 20) contains the *Bomanin* gene cluster (eight small, secreted peptides involved in immune defense) as well as *Imd* (receptor for the *Imd* pathway). Line A5 harbors two unique nonsynonymous SNPs in *Jabba* exon 6 as well as a loss of stop codon in *BomT2* (a *Bomanin*). The two *Diptericins* are tempting candidates, but neither have any variation unique to line A5 in either coding or regulatory regions. Furthermore, the DGRP lines segregate for null alleles of *DptA* and those lines are right in the middle of the line effects (51^st^ and 53^rd^ percentile). There is also a single line with a null mutation (premature stop) in *DptB* in the DGRP (line 223) and it showed 100% survival. In the DSPR, line A3 has a 37bp deletion that starts in exon 1 of *DptB* and ends in the intron, and that genotype has no apparent deficit in immune defense against *L. fusiformis*. So we favor the hypothesis that responsible variation near QTL peak 23 is in *Jabba*, one of the *Bomanins* or some other gene, but not either of the *Diptericins*.

Line effects and mapping in the DSPR A2 population were correlated when comparing Day 2 and Day 5 survival (Supplemental Figures 6A, 6B, 6F, 7, Supplemental Table S8, Pearson correlation coefficient for line effects = 0.908, Pearson correlation coefficient for LOD scores = 0.957). While the peaks overlapped significantly, the Day 2 survival did nothing to further filter to a smaller set of candidates. We therefore utilize Day 5 survival data for population A2.

We infected 391 lines from the B2 DSPR panel with a mean of 18.81 (range 10-20, median=20) individuals infected per line. The mean survival was 0.828 (median=0.85, st. dev. = 0.134). There was again considerable variation in survival across lines (Figure 4A and Supplemental Figure S6C, Supplemental Table S9). Heritability was 0.674 for survival 5 DPI and 0.760 for survival 2 DPI. The QTL scan did not result in any significant peaks with alpha = 0.05 (Figure 4B). In fact, we had to relax alpha to 0.20 to get significant peaks. There were 3 overlapping peaks on Chr3R between 15,470,000 and 16,770,000 – a region with 145 genes (qtl B5.1). This peak explained 7.35% of the variance. There are several gene s plausibly associated with immune defense. First is *Stat92E*, the transcription factor of the JAK/STAT pathway (not significantly induced, but expression was up 25.7%). Second are a cluster of *Turandots*, immune/stress effectors regulated by JAK/STAT. Of the five *Turandots* under the peak, *TotA, TotC* and *TotX* were all significantly upregulated after infection and there was no expression data for *TotB* or *TotC*. The founder genotypes under these peaks show that lines B5 and B7 have significantly lower survival than the other lines (Supplemental Figure 8C). Scanning through variation in these genes, we did not find any sites limited to B5 and B7 that were immediately good candidates for being causal. For panel B2, the correlation between LOD scores for Day 2 and Day 5 was lower and what we observed in population A (Supplemental Table S9 and S10). In fact, there was an additional peak at Day 2 (qtl B2.1, Supplemental Figure 9). This peak is only significant with alpha = 0.1, and is on 3L at 2,880,000 (LOD score = 7.57, 2 LOD drop range: 2,850,000 to 3,100,000, 8.53% of variance explained). Founder line B3 genotypes had a significantly lower survival than others (Supplemental Figure 7D). The peak includes 27 genes including *spz-5* (a paralog of the canonical *Toll* pathway *Spaetzle* gene). These genes were not differentially expressed in the transcriptome data.

### Defense against L. fusiformis shares a similar genetic architecture with defense against a fungal pathogen

We reanalyzed the data from [43], where DSPR panel A1 was infected with *Beauveria bassiani* (a fungal pathogen) and mapped survival proportion. We performed the same logistic regression (described above), which is different from their approach, and used other variables (all random effects) as they described. Our heritability estimate was 0.921 which is much higher than what they estimated (0.53). We cannot examine correlations among lines for immune defense against *B. bassiani* and *L. fusiformis* because the experiments were done in different lines (A1 vs. A2). However, we can determine whether there is significant overlap in QTLs, which would suggest a shared architecture for immune defense. We found considerable variation in the random effects of line for survival after *B. bassiana* infection (Supplemental Figure 6E, Supplementary Table S11). In contrast to what Shahrestani *et al. [43]* reported, we did find one peak significantly associated with survival after *B. bassiana* infection at alpha = 0.05 (Figure 5A, Supplemental Table S12). The peak centered at chromosome 2R:14,660,000 (Figure 5B, qtl Bbas.1, 2 LOD drop range 14,590,000-14,700,000, LOD score = 9.06, 13.15% of variance explained). The A5 founder genotype had significantly lower survival than other lines (Figure 5C).

**Figure 5.**
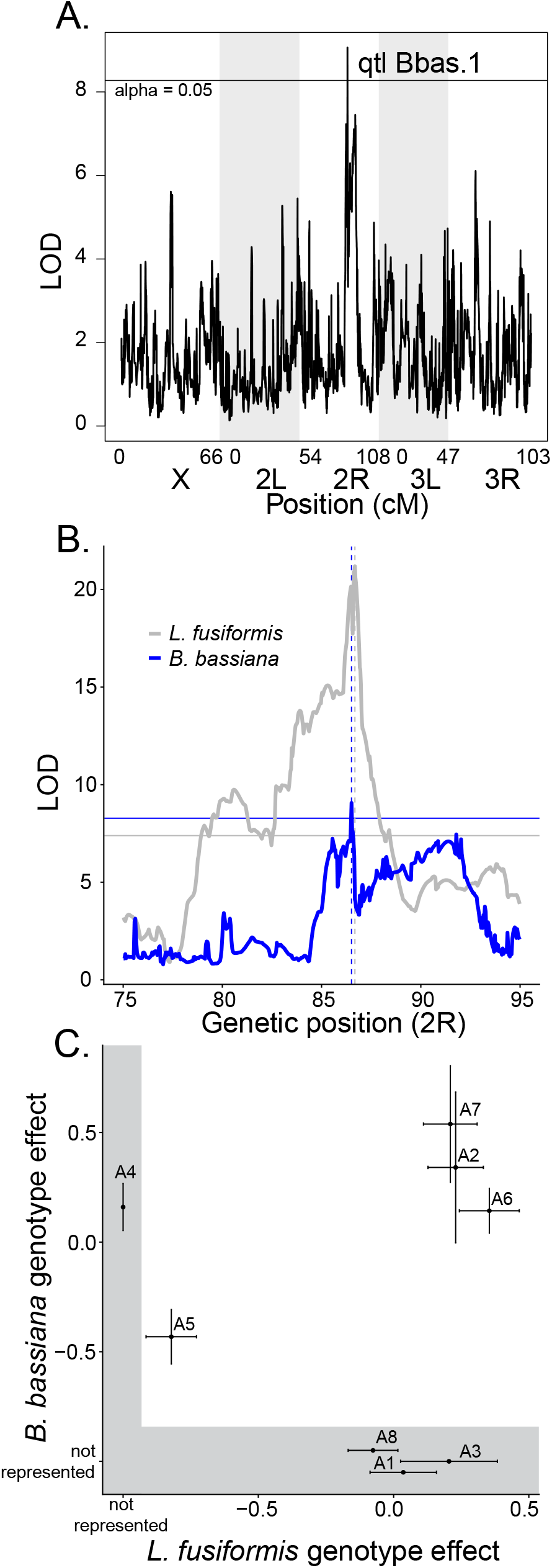
A shared genetic architecture of immune defense between *L. fusiformis* and *B. bassiana*. A) *B. bassiana* QTL analysis yields a single peak. B) The peaks on 2R for *B. bassiana* (blue, qtl Bbas.1) and *L. fusiformis* (gray, qtl A5.2) overlap. Horizontal lines represent significance threshold (alpha = 0.05), dashed vertical lines show the locations of the highest peaks. C) Founder genotype effects (plus standard error) for both experiments. Note that founder genotypes not present at that locus in one experiment are over a gray background.

Thus, the *B. bassiana* peak is quite similar to the location of qtl A5.2 for *L. fusiformis*, even though the two experiments were performed in different subsets of the A population (Figure 5B). To test whether this overlap is more than expected by chance, we performed a Fisher’s Exact Test with the fraction of physical positions with LOD score greater than the threshold score (alpha = 0.05). Day 5 survival for *L. fusiformis* had 289 of 11768 significant, while *B. bassiana* had a single position significant of 11768. That one position significant for *B. bassiana* was also significant for *L. fusiformis*, which is a significant excess over the expectation (expected overlap = 0.0246, Fisher Exact Test P-value < 0.0001). Furthermore, the founder genotype effects for both phenotypes clearly show reduced survival for the A5 genotype compared to all other genotypes present (this is true whether we match the peaks as in Figure 5C or pick the most significant peak for each genotype not shown).The DSPR experiments lead us to conclude that different panels (DSPR A2 and B2), which have different founder genotypes, have different genetic architectures for immune defense against *L. fusiformis*, but in two different mapping experiments performed in two different labs with very different pathogens, there appears to be a shared genetic architecture for resistance to infection (DSPR A2 *L. fusiformis* vs. DSPR A1 for *B. bassiana*).

### Infections of RNAi knockdown flies show the potential role of individual genes in immune defense against both L. fusiformis and B. bassiana

The number of genes under the shared QTL peak in the DSPR A population are too numerous to test individually. We therefore tested as many genes in the *Bomanin* cluster as were available and the gene encoding *Jabba*. After infection with *L. fusiformis* or *B. bassiana*, both *BomS2* and *BomBC2* knockdown flies stand out as being more susceptible to infection than controls (Figure 6, Supplemental Tables S13 to S17, P<0.001 in each comparison). Other members of the *Bomanin* cluster had little to no influence on survival after infection with either pathogen. *Jabba* fatbody knockdowns also had little impact on survival after infection. Note that the two AttP40 RFP RNAi knockdowns, used as additional controls, showed intermediate survival. Each still show significantly higher survival compared to BomBc2 (P<0.001) for both *L. fusiformis* and *B. bassiana*.

**Figure 6.**
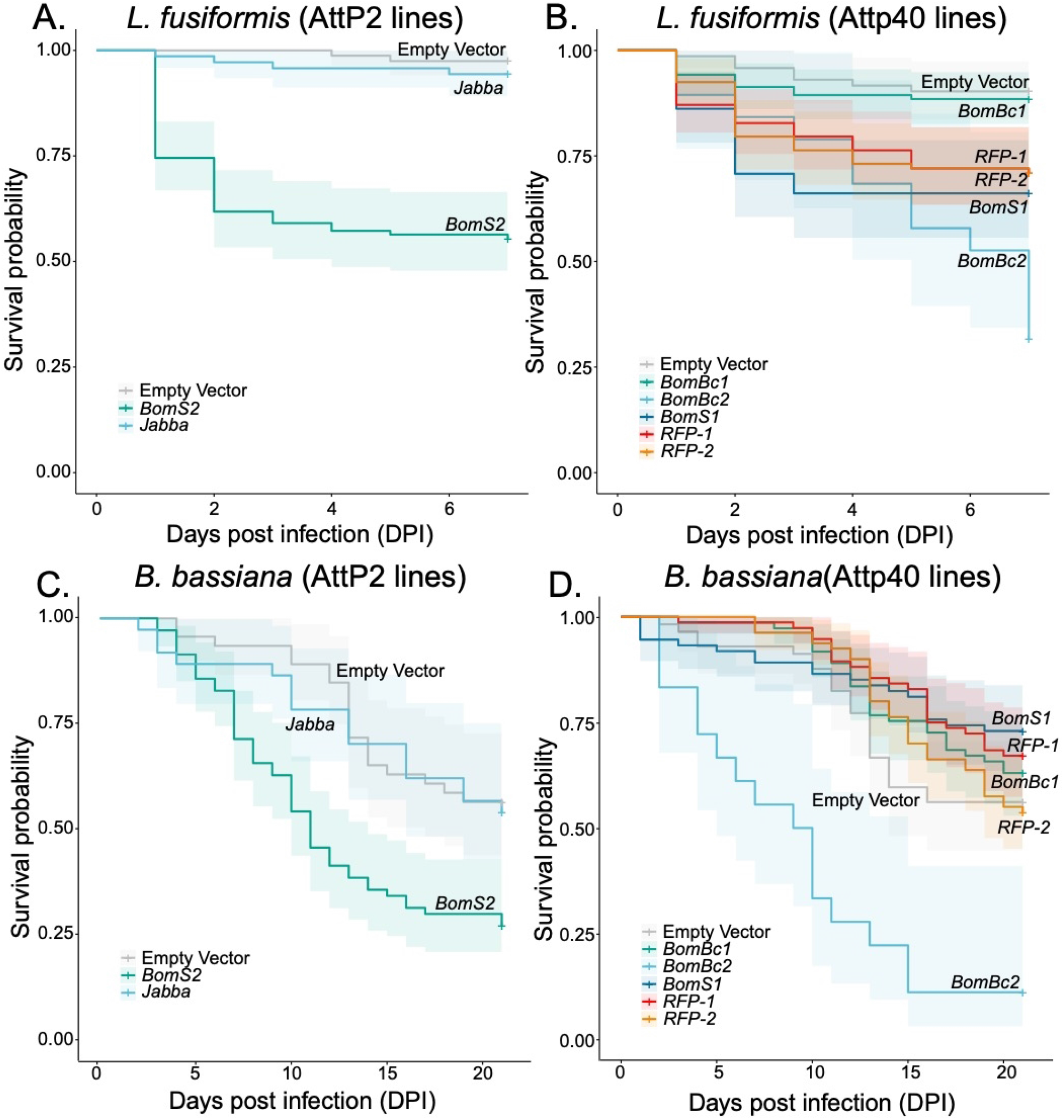
RNAi knockdowns reveal roles of specific *Bomanin* genes in immune defense against both *L. fusiformis* (A and B) and *B. bassiana* (C and D). In all cases, empty vector controls are in gray, other controls are in reddish tones and knockdowns of genes of interest are in blue/green tones.

## Discussion

Populations face infection challenge from innumerable different pathogens. The genetic factors that influence susceptibility to infection may vary from population to population for a specific pathogen, and from pathogen to pathogen within a specific host population. We examined genetic variation for susceptibility to the Gram-positive bacterium, *L. fusiformis* infection in 3 *Drosophila* mapping populations and find no evidence that the same genetic factors contribute to phenotypic variation in susceptibility across the three populations. We also compared the genetic causes of phenotypic variation for susceptibility to *L. fusiformis* and the fungal pathogen, *B. bassiana* [43]. Surprisingly, even though these experiments were conducted by different researchers in different labs with different subpopulations, there was striking concordance of a single quantitative trait locus (QTL) for both pathogens. Although that QTL peak was broad, it contains a cluster of immune peptides, some of which are induced upon infection, and show increased susceptibility to both infections upon RNAi knockdown.

We hoped to determine the extent to which different genotypes are correlated in how they defend against infection. Are some genotypes universally more resistant to infection? Is correlation stronger when pathogens are more closely related? Do correlations suggest tradeoffs among immune phenotypes (pathogen X vs. pathogen Y) or between immune phenotypes and other life history traits? The DGRP lines are widely used for measuring phenotypic variation and attempting to map the genetic causes of that variation. As such, there are several studies on both immune and life history traits to examine genotypic correlations. Although we did find evidence of correlation, it was difficult to tease apart whether these relationships were truly biological or were artifacts – stronger correlations tended to be between phenotypes measured in the same lab or institution (different pathogens in the same lab, the same phenotype for each sex). In fact, there were no significant negative correlations, which would be expected with strong tradeoffs. The only exception to the general lack of signal (positive or negative) was the positive correlation between male starvation stress resistance and female *P. aeruginosa* survival. Although this might suggest that some genotypes are better at responding to general stress (starvation or infection) than others, it is curious that the correlation is only significant for one infection type and not even in males infected with *P. aeruginosa*. We therefore hesitate to draw conclusions from this correlation.

In contrast to the DGRP experiments, we find a striking shared QTL peak comparing infection with both *L. fusiformis* and *B. bassiana* in the DSPR lines. We are unable to measure the correlation between genotypes since the two experiments were performed in two different DSPR subpopulations. Note that both subpopulations were initiated from the same set of 8 founder genotypes, so the genetic variation present in the two subpopulations is similar. The shared peak for two very different pathogens could mean several things. First, there are many genes under that peak and even many paralogs of the genes encoding *Bomanin* peptides. Therefore, it is possible that the genetic basis of increased susceptibility to *L. fusiformis* in the A5 background is different from the genetic basis of increased susceptibility to *B. bassiana* in the A5 background. If the phenotypic variation for the two pathogens was caused by different, but unique genetic changes, we imagine that natural selection might act on these linked variants as a single unit. Second, there may be a shared genetic basis for resistance to infection. Perhaps a single *Bomanin* (or small subset of *Bomanins*) are responsible for most of the response to infection, but in a nonspecific way. At this point, we know very little about how specific *Bomanins* act in the immune system (Table 1), so whether individual *Bomanin* paralogs act generally or broadly is not clear. If single *Bomanins* or even clusters of *Bomanins* act generally, times of high pathogen pressure should select for resistant (likely more highly expressing) genotypes. Such selection would be relaxed or reversed (if *Bomanins* are costly) in the absence of infection.

Our approach enabled us to also determine whether the same genetic factors influence immune defense in different populations. We acknowledge that none of the three populations are ideal populations: DGRP are inbred lines that survived the lab environment after capture at a farmer’s market, DSPR are derived from several lines from around the world and made into a “synthetic” population. Nonetheless, there was no evidence that any of the three populations shared genetic variation in any particular region that influenced survival after *L. fusiformis* infection. This may be because pathogen pressure varied in the populations from which these lines were derived. It may also be that immune alleles tend to turn over rapidly as populations coevolve with pathogens meaning that the alleles best suited to fight infection in one population are missing from another population. Finally, the small founder populations in the DSPR may preclude anything but high frequency variants from being detected in multiple experiments.

Several recent studies have attempted to characterize the role of individual *Bomanin* genes, the entire cluster of 10 genes, or subsets of those genes (summarized in Table 1) [1-3, 26, 46]. This began with Clemmons *et al*. [26] who characterized the gene family and showed that deletion of the entire cluster rendered flies highly susceptible to infection with *E. faecalis, C. glabrata* and *F. oxysporum*, but not the Gram-negative *E. cloacae*. The authors also showed the first evidence that there might be some specificity of activity since deleting the six left paralogs of the *Bomanin* cluster resulted in high susceptibility to *E. faecalis* and F. oxysporum, but not *C. glabrata*. To investigate further, Lindsay *et al*. [2] infected flies with four *Bomanin* transgenes on a *Bomanin* cluster partial deletion with *C. glabrata*. Two transgenes (*BomS3* and *BomS5*) restored survival to wildtype levels, one transgene (*BomS6*) showed intermediate restoration of survival, and one (*BomT2)* showed no restoration of wildtype survival. As expected, the expression of these genes after infection correlates with their ability to restore wildtype survival. When expressed with the *BomS3* promoter, *BomS6* was not susceptible to *C. glabrata* infection, but *BomT2* showed no difference in susceptibility. Xu *et al*. [3] showed that *BomBc1, BomS3* and *BomS4* were significantly induced upon *A. fumigatus* infection and that, as a whole, the *Bomanin* cluster enhances both resistance to, and tolerance of, infection with *A. fumigatus. BomBc1, BomS3* and *BomS6* increased tolerance to the fungal toxin, restrictocin, whereas *Bom S6* and *BomS1* increased tolerance to another toxin, verruculogen. Here, we showed that natural variation in immune defense against both the Gram-positive *L. fusiformis* and the fungal *B. bassiana* map to a QTL that contains the *Bomanin* cluster and that knockdown of specific *Bomanin* copies result in enhanced susceptibility (note that we do not differentiate between tolerance and resistance). Unfortunately, most of these studies utilize a subset of the *Bomanins* and use different techniques (CRISPR/Cas9 knockouts, transgenic lines, RNAi), so a systematic understanding of the specificity of individual *Bomanins* or the subclasses (tailed, short and bicipital) remains elusive.

This study highlights the context-dependence of the causes of genetic variation in immune defense. Within the same species, different populations harbor different genetic variants that result in variation in immune defense. Surprisingly, however, within a population, two very different pathogens share a QTL for susceptibility to infection, with a single founder genotype being significantly more susceptible than others. Although our approach does not have the resolution to determine whether the causal genetic variants are the same or just genetically linked, this linkage allows selection to act more broadly to reduce susceptibility when susceptibility to such disparate pathogens is governed by a single locus.

## Methods

### Drosophila strains and husbandry

D*rosophila* Synthetic Population Resource (DSPR) lines from populations A1 and B1 are maintained by Stuart Macdonald at the University of Kansas [39]. The DSPR population lines were derived from two different sets of founder inbred lines. Each population was interbred among lines for several generations, then inbred to create the recombinant inbred lines used in these experiments. *Drosophila* Genetic Reference Panel (DGRP) lines were obtained from Brian Lazzaro at Cornell University and from the Bloomington *Drosophila* Stock Center (Bloomington, IN). Canton S is a standard laboratory *Drosophila* line. All *Drosophila* stocks were maintained on standard molasses media at 23 degrees Celsius on a cycle of 12 hours light to 12 hours dark. We refer to both sets of lines by their numeric assignment (for the DGRP we refer to line 321, not RAL_321 or line_321).

### Microbes and microbiology

*Lysinibacillus fusiformis* (strain Juneja) is a natural pathogen of *Drosophila* obtained from Brian Lazzaro at Cornell University [45]. Bacteria were grown from single colonies in LB liquid media overnight to stationary phase, then concentrated to an optical density (600nm) of about 4.0. *Beavaria bassiana* (strain GHA) was obtained from Parvin Shahrestani (California State University at Fullerton). Spores were grown in malt extract agar, then resuspended with a 10ml syringe, filtered through a 0.22 μm Miracell filter and resuspended in 20% glycerol at 4.3e08 spores per mL as measured on a hemocytometer.

### General Infection protocol

Flies were tipped to new media and cleared after 2-4 days of egg laying to control larval density. Ten to fourteen days later, emerging adults were moved to new media vials. Three- to seven-day old males were infected after light CO_2_ anesthesia by pricking in the thorax with a needle dipped in bacteria or fungal suspension or LB broth (or 20% glycerol) as a sterile control. The number surviving 2- and 5-days post infection was noted. For Canton S infections and RNAi knockdowns, we noted survival daily. Infections were performed over several weeks with 3-4 days of infection each week. A single researcher performed all infections (n=22174) for the GWAS and QTL experiments, and a different researcher performed all infections for the RNAi experiments. These experiments were done in batches of subsets of lines since it would not be possible to infect all lines in one batch. Each subset of lines was chosen randomly and batch effects were included in statistical models.

### Infection dynamics in Canton S

The effect of *L. fusiformis* infection on Canton S survival was assessed using a Cox Propotional Hazard test implemented in the R packages *survival* and *survminer [59, 60]* with treatment (infected vs. sterile prick) as the independent variable.

### RNA-sequencing of infection in Canton-S males

We used Canton S flies to compare expression 12 hours after infection with *L. fusiformis* to those uninfected (no sterile prick). Each treatment contained 3 replicates of 3 flies per replicate. We flash froze flies in liquid nitrogen, then extracted RNA using a Zymo Quick-RNA Microprep kit (R1050, Zymo Research, Irvine, CA). We then constructed RNA libraries using the Lexogen QuantSeq 3’ mRNA-Seq Library Prep kit (Lexogen GmbH, Vienna, Austria). We sequenced the samples on an Illumina MiSeq Micro SR100 run. Raw Reads were quality filtered using the fastx toolkit (http://hannonlab.cshl.edu/fastx_toolkit/; RRID:SCR_005534), then mapped with STAR version 2.7.10b [61, 62] to the *Drosophila melanogaster* (r6.18) genome, processed with samtools version 1.9 [63, 64] and analyzed with DESeq2 [65] using a simple infection vs. no prick design.

### Genetic variation for immune defense in the DGRP

We infected male DGRP flies with OD_600_ ∼ 4.0 *L. fusiformis*. Each vial consisted of eight to ten infected flies with one to five vials per line for the DGRP. This corresponds to an average of 36.0 flies infected per line in 199 DGRP lines.

We performed the DGRP genome wide associations with and without line 321. We used both approaches because line 321 showed unusually low survival at 5 days post infection (DPI) and others [1, 21] noted and excluded 321 in their analysis of resistance to *Metarhizium anisopliae* and *Enterococcus faecalis* infections. However, line 309 actually had lower survival (0% at 5 DPI) and we hesitate to start dropping all lines with low survival since variation is the point of this type of mapping. We measured survival both 2- and 5-days post infection and performed the mapping experiment at both time points. We fit a mixed effects logistic regression model using the *lmer* function in *lme4* [66] and line effects were extracted using the *ranef* function. These values were used with the DGRP website [30] to perform association mapping. Thus, there were four total mapping experiments for the DGRP: 2-and 5-days post infection and with and without line 321. We calculated heritabilities using line variance and total variance from the *VarrCorr* function in *lme4*.

We next asked whether line effects correlated between DGRP infection experiments (and a few other traits). We retrieved line effects for both male and female 24 hour bacterial load and female survival post *Providencia rettgeri* infection [5, 37], male and female *Coxiella burnetii* hazard ratio [67], female *Pseudomonas entomophila* enteric survival [68], male and female *Pseudomonas aeruginosa* survival [21], male and female *Metarhizium anisopliae* survival [21], *Staphylococcus aureus* phagocyte mobilization [69], *Listeria monocytogenes* bacterial load [70], *Enterococcus faecalis* survival [1], male aggression [71], male starvation resistance [30], pigmentation (tergite 5) [72], lifespan and fecundity [73], and weight [34]. We calculated Spearman’s rank correlations for these 20 phenotypes and grouped by non-immune, Gram-positive/fungal and Gram-negative pathogens. In these analyses, we used the negative of the load values which means that all immune phenotypes have higher values indicating better immune defense and lower values meaning worse immune defense.

### Genetic variation for immune defense in the DSPR

The DSPR experiment was similar to that for the DGRP, but with more lines and fewer individuals infected per line. We used from one to four vials, consisting of 8 to 10 individuals, per line in the DSPR population. This corresponds to an average of 22.3 flies infected per line in 343 lines from DSPR population A2 and 18.8 flies infected per line in 391 lines from DSPR population B2. For each population, we used a logistic regression for proportion alive at day 2 and day 5 with line and infection date as random effects. These random effects of line were used directly in the DSPRqtl package in R [39, 40]. We used the DSPRscan function with the model effect∼1. To determine a significance threshold, we performed 1000 permutations using the same model. Peaks were determined using the LOD drop 2 method in the DSPRPeaks function and nearby and overlapping peaks were curated by hand.

We also retrieved raw data for survival after *B. bassiana* infection [43] and performed a logistic regression analysis with random effects of line, replicate nested in line, group and group nested in set (as in their analysis). This differs from the authors’ approach who used arcsin-square root transformed survival. All other analyses were performed as above.

### RNAi knockdown experiments to examine the role of individual genes in immune defense

We followed up our QTL mapping by examining several genes that occur under the main peak on chromosome 2R. We obtained the TRIP RNAi lines [74] from the Bloomington Drosophila Stock Center (Bloomington, Indiana, USA) and crossed to the C564 (RRID: BDSC_6982) fat body-specific driver [75]. We also crossed the driver to an empty vector and RFP RNAi (when available) as negative controls. Infections were performed as described above for *L. fusiformis*, but *B. bassiana* were monitored for 21 days instead of 7 days, with flies transferred to new food every 3-4 days. In addition, we mock infected lines with either Luria-Bertani broth or 20% glycerol as a sterile wound control. Very few of these flies died from the wound. We assessed significant differences in survival using the *coxph* function from the *survival* package in R [59, 76] with genotype and date as variables and the empty vector as the reference. Since the RNAi lines are on different backgrounds (AttP2 and AttP40), we performed separate analyses based on background.

## Acknowledgements

We thank Brian Lazzaro for providing the *L. fusiformis* strain, Stuart Macdonald for the DSPR lines and advice on mapping, and Unckless lab members for helpful comments and advice. The Bloomington Drosophila Stock Center provided several DGRP and RNAi lines. RNA sequencing was performed in the KU Genome Sequencing Center, supported by the National Institute of General Medical Sciences of the National Institutes of Health under award number P30-GM145499. RLU was supported by NIH grant R01-AI139154.

## Supplemental Tables (See Excel File)

Table S1. Gene Ontology enrichment for expression 12 hours after *L. fusiformis* infection in *D. melanogaster* Canton S males.

Table S2. Values for DGRP random effects from the logistic regression model. All lines included or line 321 removed and at 5- or 2-days post infection.

Table S3. SNPs significantly associated with survival 5 days post infection in the DGRP with all lines.

Table S4. SNPs significantly associated with survival 5 days post infection in the DGRP when line 321 is omitted.

Table S5. SNPs significantly associated with survival 2 days post infection in the DGRP with all lines.

Table S6. SNPs significantly associated with survival 2 days post infection in the DGRP when line 321 is omitted.

Table S7. Values for DSPR random effects from the logistic regression model for survival 5 days post infection with *L. fusiformis* in population A.

Table S8. Values for DSPR random effects from the logistic regression model for survival 2 days post infection with *L. fusiformis* in population A.

Table S9. Values for DSPR random effects from the logistic regression model for survival 5 days post infection with *L. fusiformis* in population B.

Table S10. Values for DSPR random effects from the logistic regression model for survival 2 days post infection with *L. fusiformis* in population B.

Table S11. Values for DSPR random effects from the logistic regression model for survival post infection with *B. bassiana* in population A.

Table S12. LOD scores for all genomic positions for all 5 mapping experiments (populations A2 and B2 for 2- and 5-days post infection for *L. fusiformis* and A1 for *B. bassiana)* in the DSPR.

Table S13. Raw data for RNAi infection with date dead and status (0 if alive at the end of the experiment).

Table S14. Genotype and block effects for survival analysis after infection with *L. fusiformis* for RNAi knockdown lines on the AttP2 background.

Table S15. Genotype and block effects for survival analysis after infection with *L. fusiformis* for RNAi knockdown lines on the AttP40 background.

Table S16. Genotype and block effects for survival analysis after infection with *B. bassiana* for RNAi knockdown lines on the AttP2 background.

Table S17. Genotype and block effects for survival analysis after infection with *B. bassiana* for RNAi knockdown lines on the AttP40 background.

## Supplemental Figures

**Figure S1.**
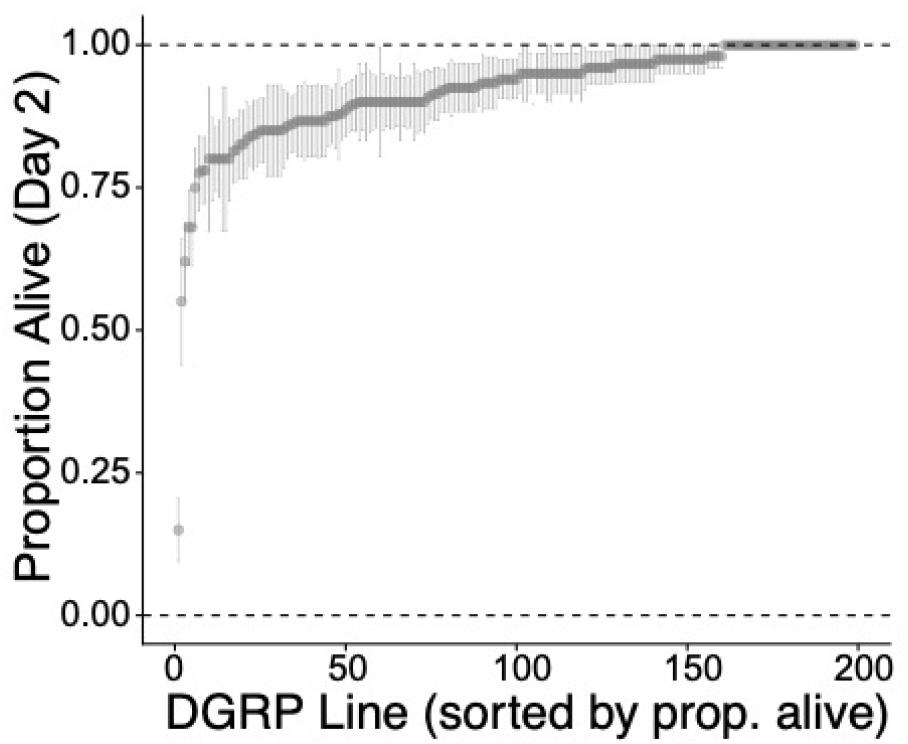
DGRP Day 2 raw survival with standard error of the proportion. Survival measured as the proportion alive at Day 2 divided by the total number infected per line.

**Figure S2.**
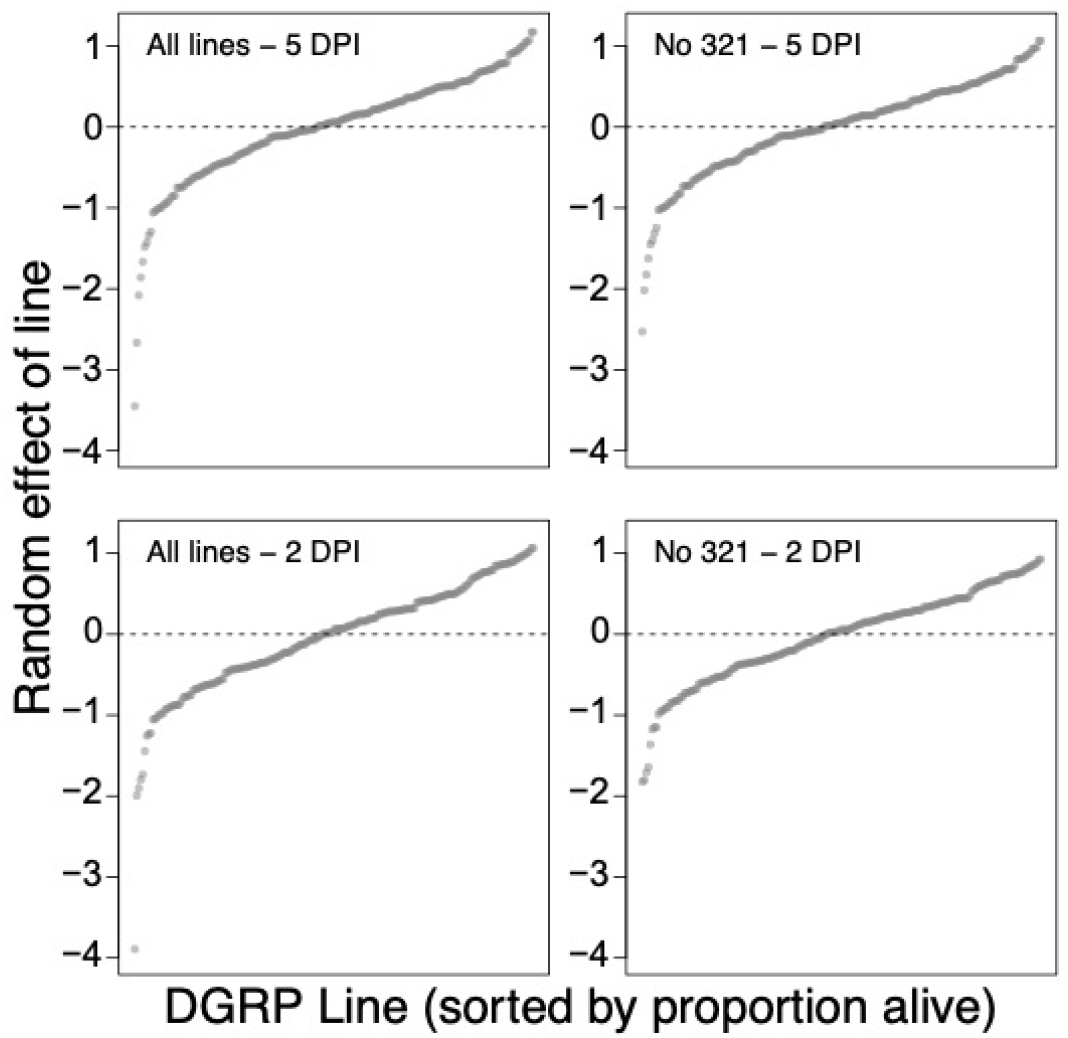
DGRP Random effects sorted by effect size from mixed effects logistic regression model. Data is presented with all lines present or excluding line 321 (which had low survival) and at 5- or 2-days post infection.

**Figure S3.**
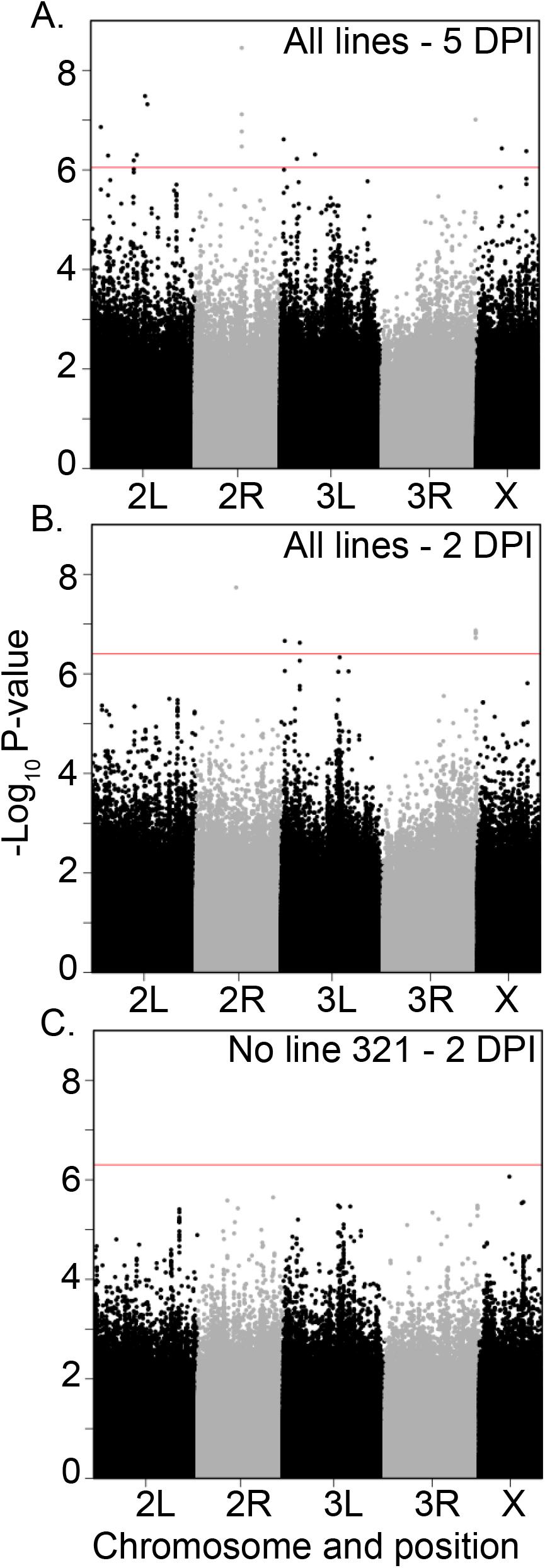
Manhattan Plots for DGRP Genome Wide Association Studies at 5- or 2-days post infection and with or without line 321 included.

**Figure S4.**
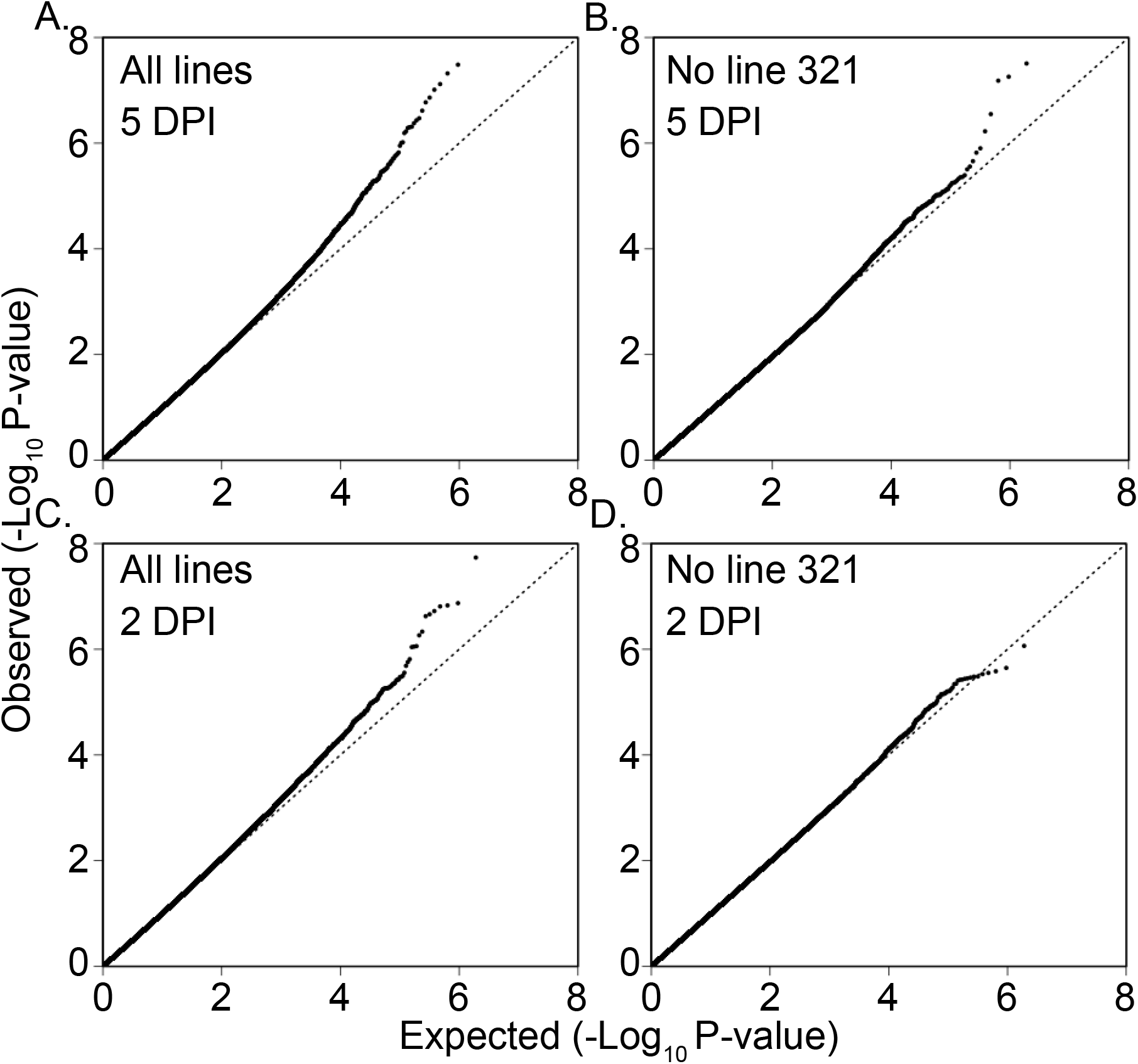
QQ Plots for DGRP Mapping Experiments at 5- and 2-days post infection and with all lines or with line 321 excluded.

**Figure S5.**
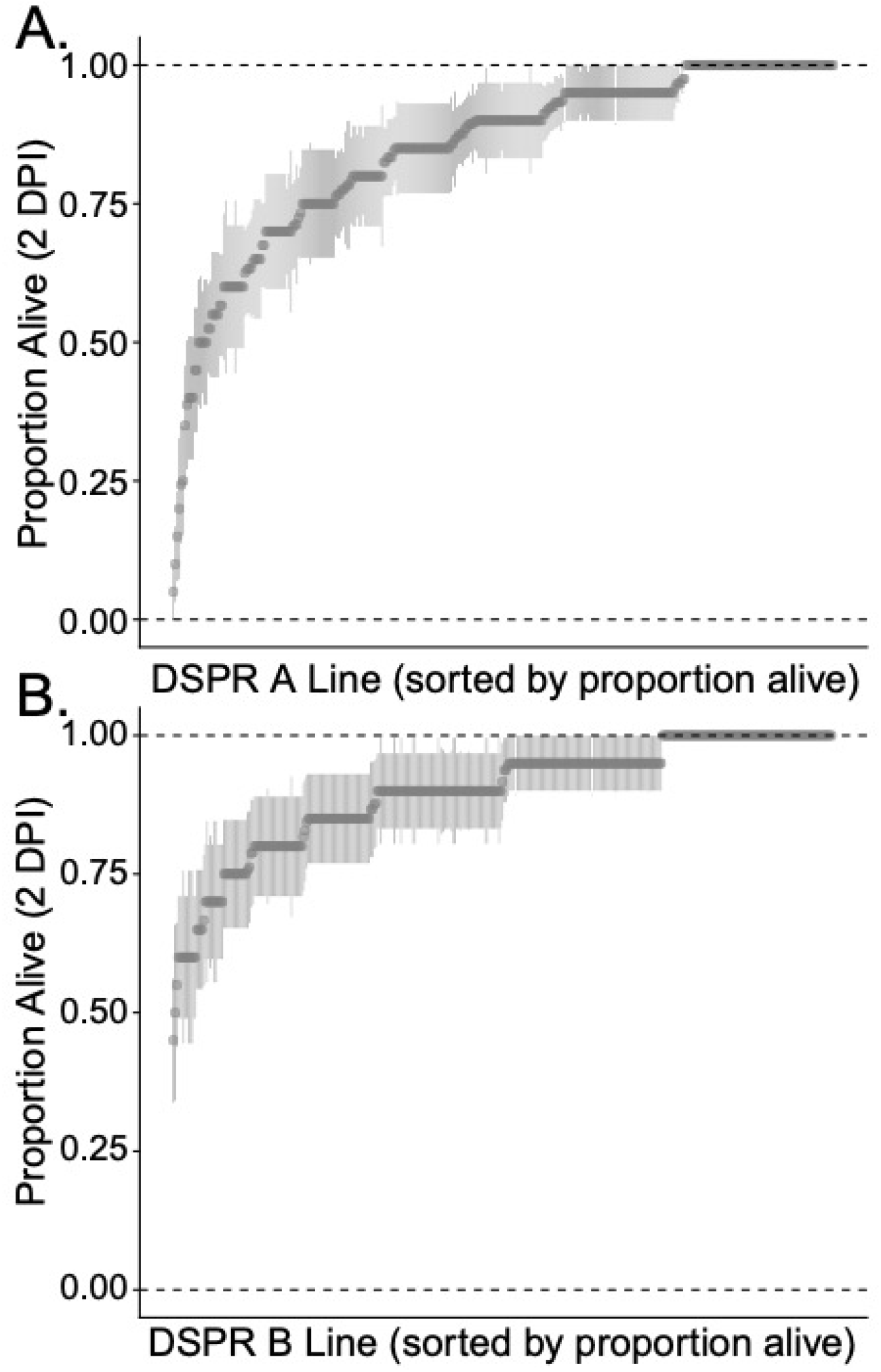
DSPR Day 2 Raw Survival plus standard error of the Proportion. Raw survival is measured as proportion alive at 2 days post infection. A) Panel A, B) Panel B.

**Figure S6.**
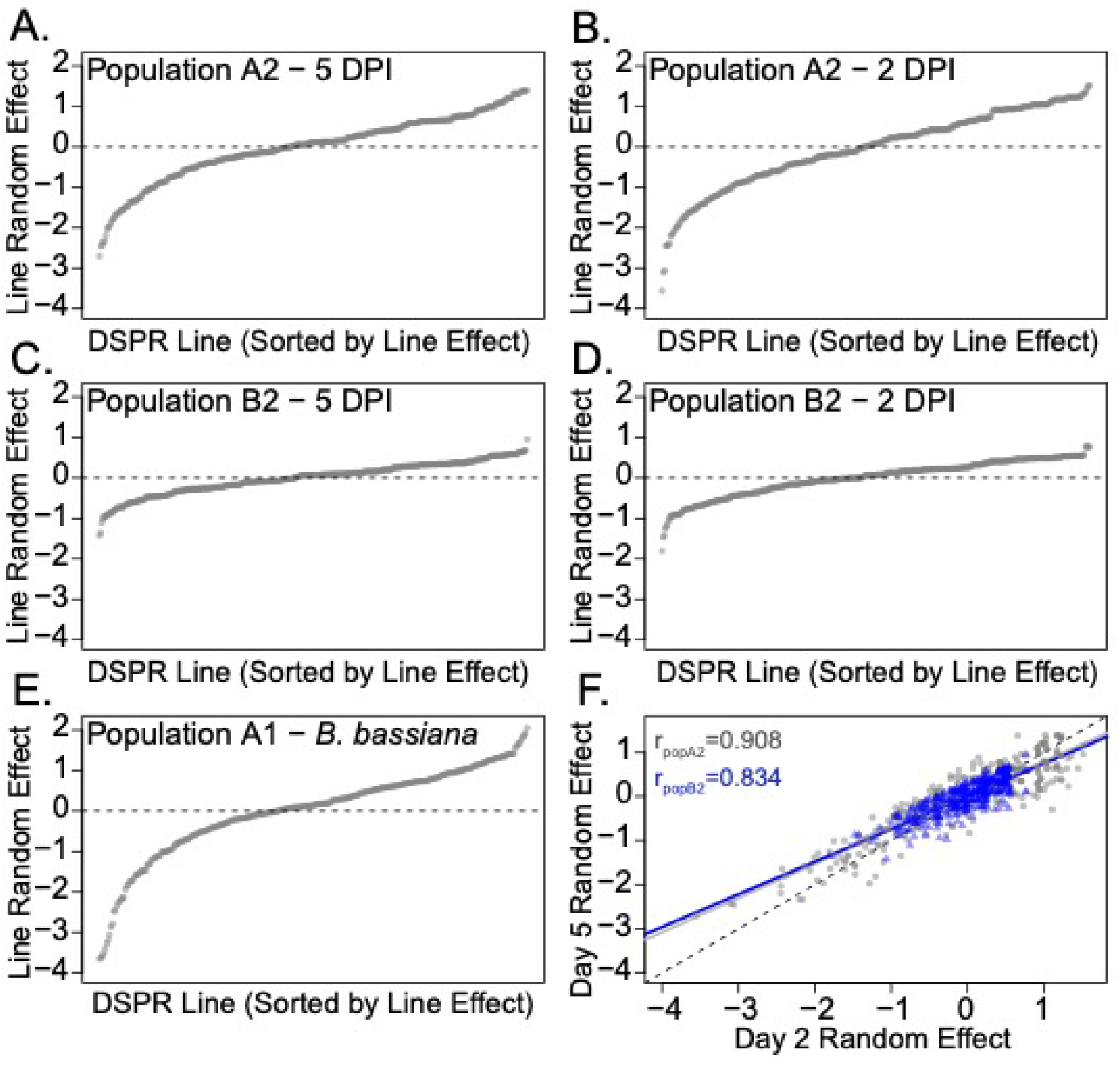
DSPR Line effects for (A-D) *L. fusiformis* survival day 2 or day 5 post infection in populations A2 and B2, E) *B. bassiana* and F) correlation between Day 5 and Day 2 line random effects in population A2 (gray) and B2 (blue).

**Figure S7.**
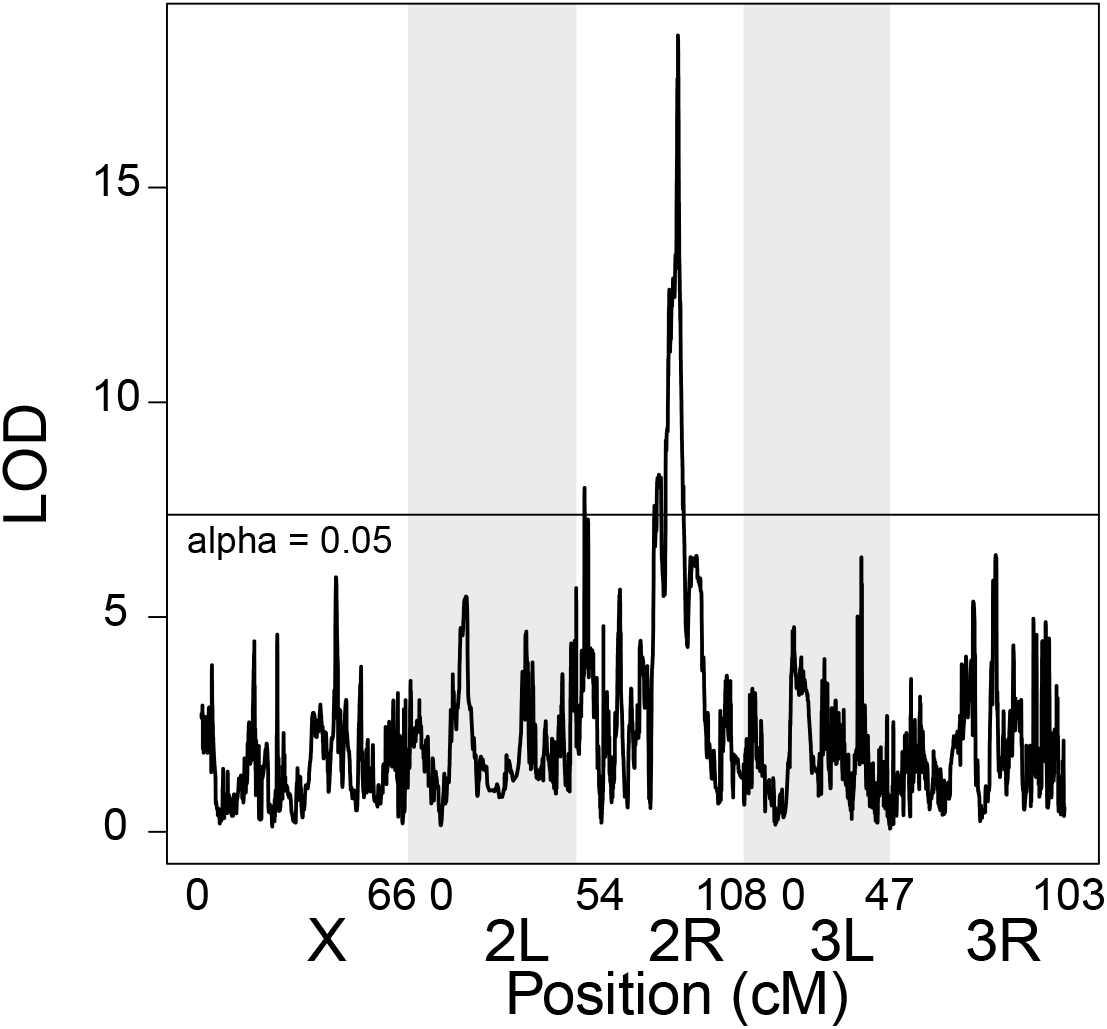
DSPR Scan plot for Day 2 survival random effect in DSPR population A2.

**Figure S8.**
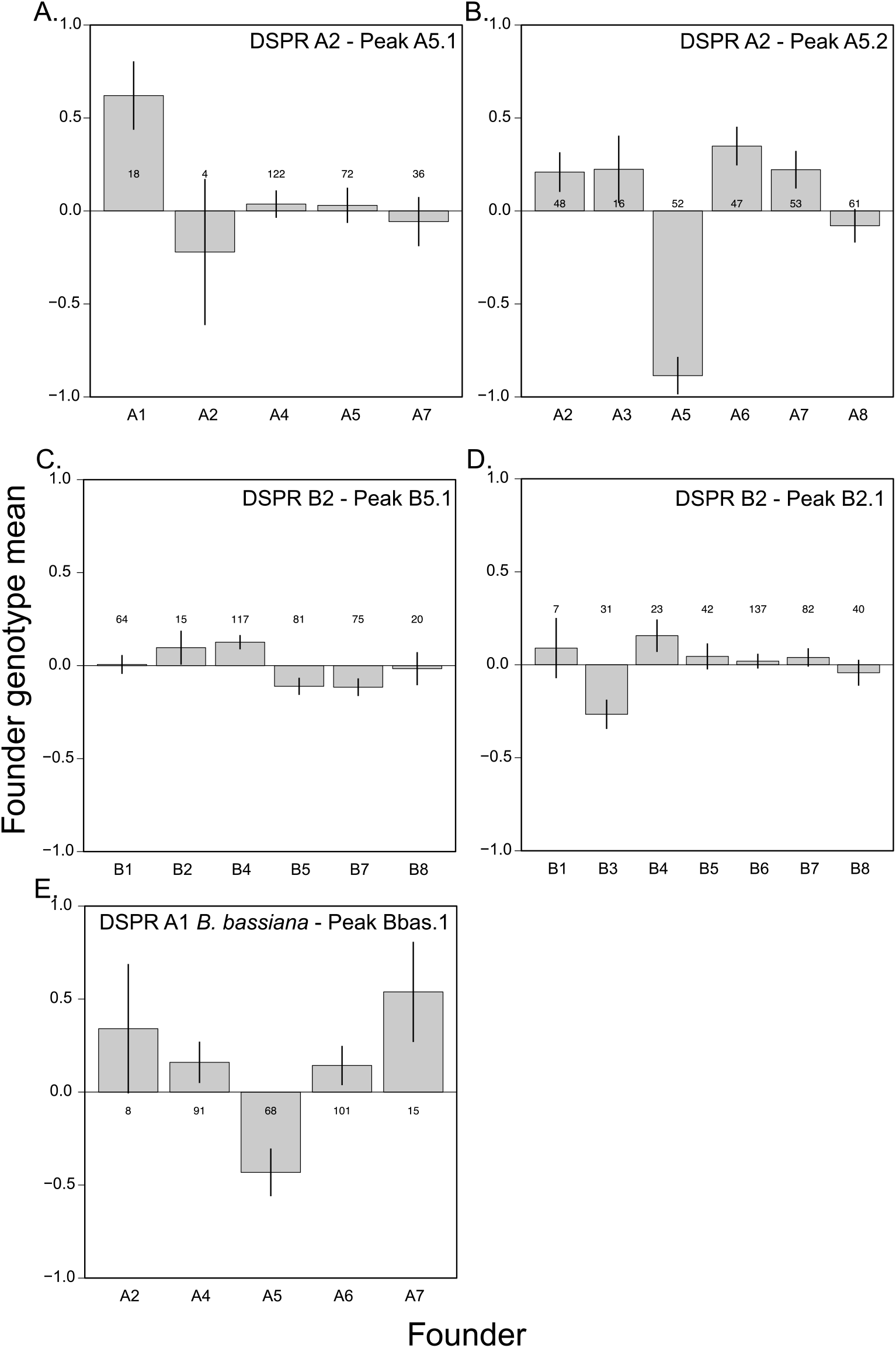
Founder genotype effects for significant QTL peaks in DSPR populations. (A and B) Population A2 Day 5 peaks 1 and 2, C) Population B2 Day 5 peak 1, D) Population B2 Day 2 peak 1, E) Population A1 *B. bassiana* survival peak 1.

**Figure S9.**
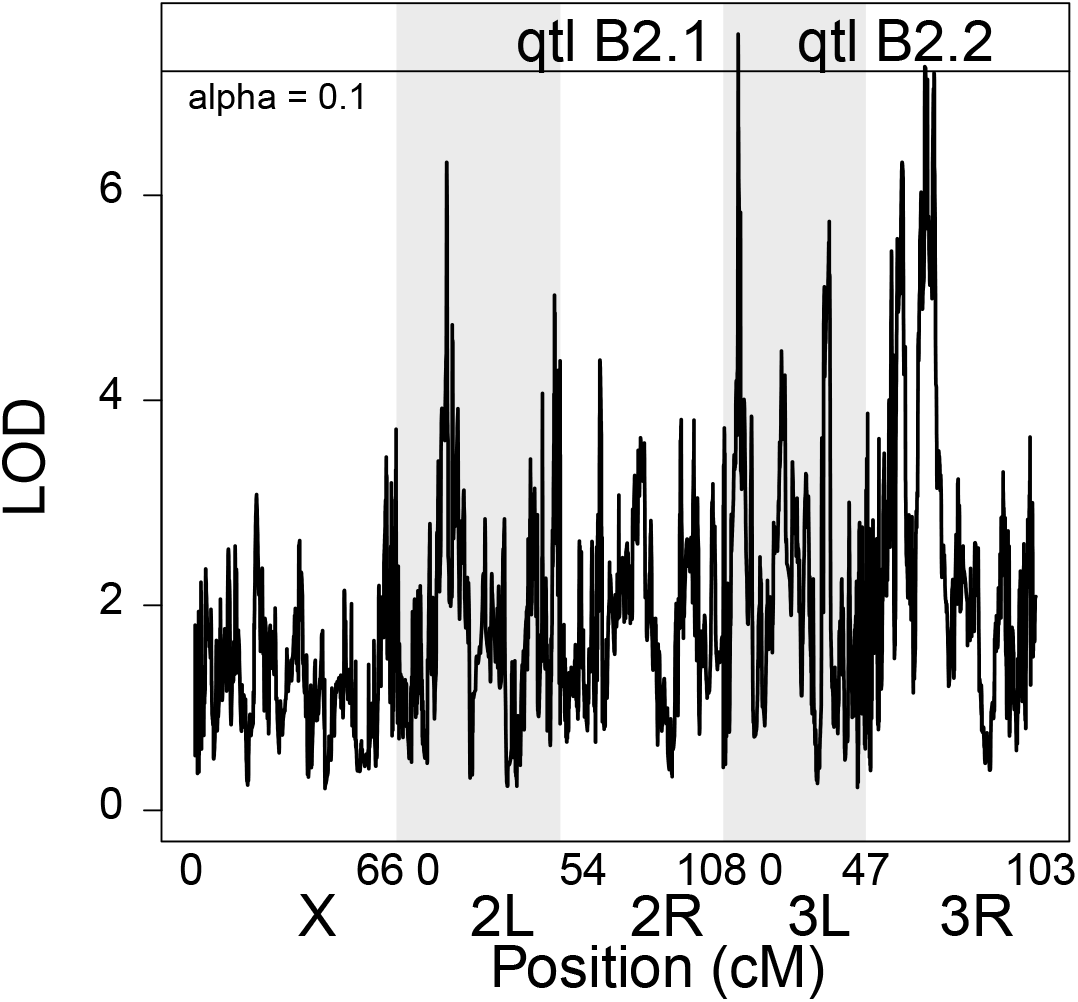
DSPR Scan plot for Day 2 survival random effect in DSPR population B2.

